# How subtle changes in 3D structure can create large changes in transcription

**DOI:** 10.1101/2020.10.22.351395

**Authors:** Jordan Xiao, Antonina Hafner, Alistair N. Boettiger

## Abstract

Animal genomes are organized into topologically associated domains (TADs), which exhibit more intra-domain than inter-domain contact. However, the absolute difference in contact is usually no more than twofold, even though disruptions to TAD boundaries can change gene expression by 8-10 fold. Existing models fail to explain this superlinear transcriptional response to changes in genomic contact. Here, we propose a futile cycle model where an enzyme stimulated by association with its products can exhibit bistability and hysteresis, allowing a small increase in enhancer-promoter contact to produce a large change in expression *without* obvious correlation between E-P contact and promoter activity. Through mathematical analysis and stochastic simulation, we show that this system can create an illusion of enhancer-promoter specificity and explain the importance of weak TAD boundaries. It also offers a mechanism to reconcile recent global cohesin loop disruption and TAD boundary deletion experiments. We discuss the model in the context of these recent controversial experiments. Together, these analyses advance our interpretation and understanding of cis-regulatory contacts in controlling gene expression, and suggest new experimental directions.

## Introduction

The genomes of many organisms have been shown to adopt a domain-like structure commonly referred to as topologically associated domains or TADs (Jerković et al., 2020; McCord et al., 2020; Rowley and Corces, 2018), defined as contiguous regions of the genome where intra-region proximity is greater than inter-region (Dixon et al., 2012; Nora et al., 2012). When plotted as a heat map of contact frequency as a function of two genomic coordinates, TADs appear as boxes on the diagonal at specific genomic coordinates. They can be detected by a range of distinct techniques, including methods relying on proximity ligation like 3C/Hi-C (Jerković et al., 2020; McCord et al., 2020; Rowley and Corces, 2018), ligation-free sequencing methods like GAM (Beagrie et al., 2017) and SPRITE (Quinodoz et al., 2018), microscopy methods such as ORCA (Bintu et al., 2018; Mateo et al., 2019), and even live cell measurements like DAMC (Redolfi et al., 2019). Recent microscopy experiments have shown TADs are a statistical property that emerges from a population of cells dynamically exploring multiple conformational states, rather than a static structure found in all cells (Bintu et al., 2018).

Despite broad agreement on the existence of TADs, their role in transcriptional regulation has recently become a subject of intense controversy, as different recent results have been interpreted to support or refute a functional link (Finn and Misteli, 2019a; Ghavi-Helm et al., 2019; Mir et al., 2019). For example, many TAD boundaries demarcate regions of co-expressed genes and separate differentially expressed genes, suggesting they play a role in cs-regulatory specificity (Long et al., 2016; McCord et al., 2020; Spielmann et al., 2018). Opposing this interpretation, it has been noted that the quantitative difference in interaction frequency is generally small: inter-TAD contacts often reach only half the frequency of intra-TAD contacts, raising questions of how such small differences could explain such clear separation of expression (McCord et al., 2020). The disruption of TAD boundaries by cis-mutation has been proposed as the causal driver of the substantial alterations in gene expression for a variety of disease conditions, a conclusion supported by animal models (Dowen et al., 2014; Franke et al., 2016; Hnisz et al., 2016; Lupiáñez et al., 2015) reviewed in (Long et al., 2016; McCord et al., 2020; Spielmann et al., 2018). However, global disruption of TAD boundaries by depletion of boundary associated factors led to few substantial changes in gene expression in cell culture models (Nora et al., 2017; Rao et al., 2017; Schwarzer et al., 2017), contradicting the earlier studies. Genetic experiments have made it clear that enhancers act on promoters only *in cis,* suggesting a requirement for physical proximity that would explain the functional role of TADs (Furlong and Levine, 2018). However, recent microscopy experiments uncovered little correlation between enhancer-promoter (E-P) proximity and nascent transcription activity at well-studied loci (Alexander et al., 2019; Mateo et al., 2019) reviewed in (Finn and Misteli, 2019b). This surprising disagreement has led to various speculative explanations, such as action at a distance through condensates with diameters tens of molecules across, or regulation lag in enhancer control (Alexander et al., 2019; Heist et al., 2019).

We wondered whether these apparent contradictions can be reasonably reconciled with known biochemical mechanisms. To answer these questions we examined simple biophysical models of proximity-dependent E-P communication grounded in known physical laws (chemical master equation). We identify mechanisms whose relevance for the spatial organization of the genome has not previously been considered, which we show provide a simple reconciliation of all three apparent contradictions. We discuss the implications for this revised view of contact-mediated cis-regulation of gene expression for interpreting experimental results.

## Results

### A simple model of hypersensitivity to changes in contact frequency

To better understand the connection between E-P contact frequency and transcriptional behavior, we started with a quantitative analysis of recently published super-resolution microscopy data (Mateo et al., 2019). These experiments used Optical Reconstruction of Chromatin Architecture to observe 3D path of chromatin through two neighboring regulatory domains. The upstream domain contained the gene *Ubx* and its enhancers, and the downstream region contained the gene *abd-A* and its enhancers (**Fig. 1A**). In wildtype embryos, interactions between the two domains, including enhancers and promoters, were less frequent than within the domains, producing two clearly distinct TADs in population average data (**Fig 1A**). Deletion of a few kilobase elements at the border of these TADs resulted in a clear increase in the interaction between the domains, though only by a factor of 1-2.5 fold across the population (**Fig 1A**). While the structures are easily distinguished at the population level, the structure of single cells are more variable, with 25% of individual wildtype cells showing some cross-TAD border mixing between the *Ubx* enhancers and the *abd-A* promoter, compared to 50% of mutant cells (**Supp. Fig 1A**). The change in gene expression, as assayed by smFISH, is hypersensitive to this moderate change in contact frequency, with an average 5-fold increase in mRNA counts of *abd-A* (**Fig. 1B).** Similar changes are also seen for *Abd-B* (**Supp. Fig. 2**). From these single cell-analyses, we concluded that the quantitative disconnect between weak TAD borders (see also **Supp. Fig 3**) and the large transcriptional effects of border disruption (see also **Supp. Fig. 4**) is not an artefact of Hi-C or of population averaging. Instead, small differences in the frequency of contact appear sufficient to drive large changes in expression, requiring us to reexamine the text-book picture of E-P regulation (Alberts et al., 2018).

**Figure 1.**
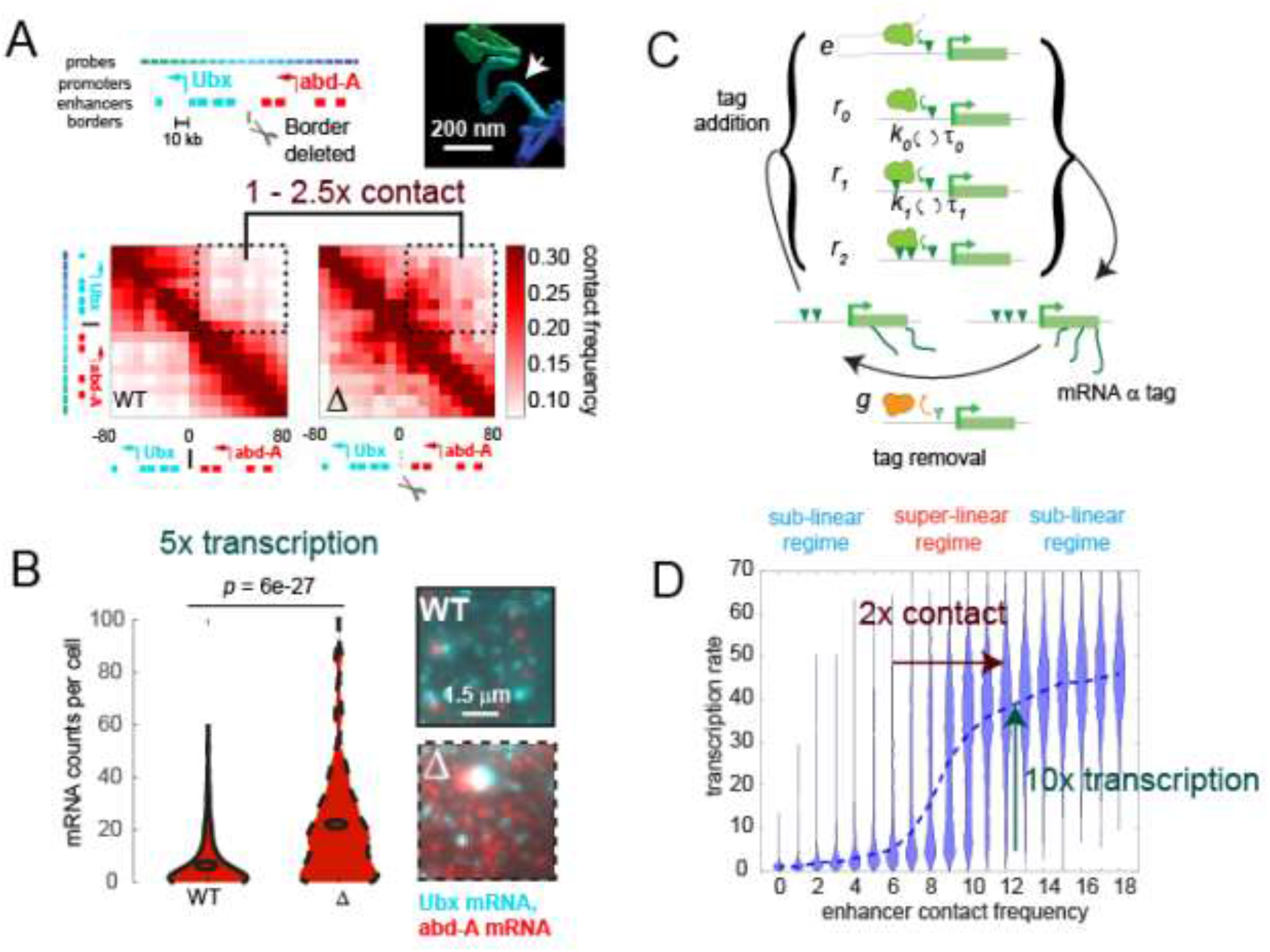
A futile cycle promoter explains super-linear response in transcription following subtle changes in enhancer promoter contact. **(A)** Quantification of the minor difference in cross border contact frequency between the *Ubx* and *abd-A* regulatory domains in wildtype and domain-border-deleted embryos, quantified by Optical Reconstruction of Chromatin Architecture, reproduced from (Mateo et al., 2019). **(B)** Quantification of the major change in transcription of *abd-A* between wildtype and domain-border deleted embryos. Raw data taken from (Mateo et al., 2019) quantified here. **(C)** Schematic of the chemical states and transition rates in the futile cycle model of enhancer-mediated promoter activity. Green inverted triangles denote promoter ‘tags’. See text for details. **(D)** Violin plot of the distribution of transcriptional activity observed for different E-P contact rates, from a simulation results from the model in C (parameters values in supplement). Dashed blue line connects the population median transcription rates. The distinct model regimes are labeled above.

Inspired by the hypersensitive cell-signaling pathways dependent on futile-cycle competition between phosphatases and kinases, we considered a model of enhancer-mediated transcription (**Fig. 1C**) in which the transcription rate of the promoter is governed by the amount of transcription promoting ‘tag’ that accumulates there. For simplicity, we assume transcription is linearly proportional to tag amount. The tag could be an accumulating condensate of transcription factor, or a histone modification, such as acetylation. The promoter has an intrinsic rate of tag addition (*r*_0_) and removal (*g*), and the enhancer regulates this competition by contributing to tag addition when it interacts (*e*) with the promoter (**Fig. 1C**). Finally, we assume the enzyme that adds the tag can also physically interact with this output product, increasing its effective addition rate from *r*_0_ to *r_0_*, *r_1_*, *r_2_* …, *r_n-max_* where *n* is the number of tag molecules currently associated, and *r_n_* > *r*_*n*-1_. This could equivalently be achieved by stabilizing the binding of enzyme by associating with the tag, increasing its local concentration instead of catalytic rate. The probabilities of product association and dissociation are proportional to the amount of existing product, with respective rate constants *k_n_* and *τ_n_*. The importance of this product interaction, the constraints on the parameter values, and range of known molecular factors which may execute these behaviors, are discussed below.

This model readily exhibited a hypersensitive regime in which small fold changes in E-P contact lead to large fold changes in transcription (**Fig. 1D**), around the experimentally observed range (**Fig 1A, 1B**). This hypersensitive behavior suggests that the relatively weak effects of most TAD boundaries (**Supp. Fig. 3**) may still have important consequences for the control of gene expression.

### Futile-cycle promoters can create an illusion of enhancer-promoter specificity

We next explored how differences in promoter properties affect enhancer-promoter responsiveness. We considered two promoters which both experience a two-fold increase in enhancer-promoter interaction at the start of the simulation (**Fig. 2A**), for example, due to removal of a neighboring strong TAD boundary. Both promoters start in a low transcription state, but the second one has a slight (two-fold) higher intrinsic tag removal rate, *g* (**Fig. 2A**). Promoter 1 exhibited a large increase in transcription, yet Promoter 2 remains almost unaffected by this perturbation, exhibiting only a small shift (**Fig. 2B**). Because of its higher intrinsic affinity for the tag removal machinery, Promoter 2 is still in the sub-linear response regime (**Fig. 1D**). Further increase in enhancer promoter interaction would be sufficient to drive it into a highly transcribing state like Promoter 1. This behavior is interesting, as many experimentally reported perturbations involve bystander genes that respond little to gains in enhancer activity (Ghavi-Helm et al., 2019; Guo et al., 2015; Paliou et al., 2019; Williamson et al., 2019), a behavior which has been interpreted as evidence of a “lock-and-key” mechanism for gene regulation (van Arensbergen et al., 2014). In that model, enhancer-promoter specificity is achieved by different distinct pairs of biochemical locks and keys, rather than by spatial organization (**Fig. 2C**). These simulations illustrate that the apparent specificity may be an illusion resulting from different tipping points in the type of promoters considered here.

**Figure 2.**
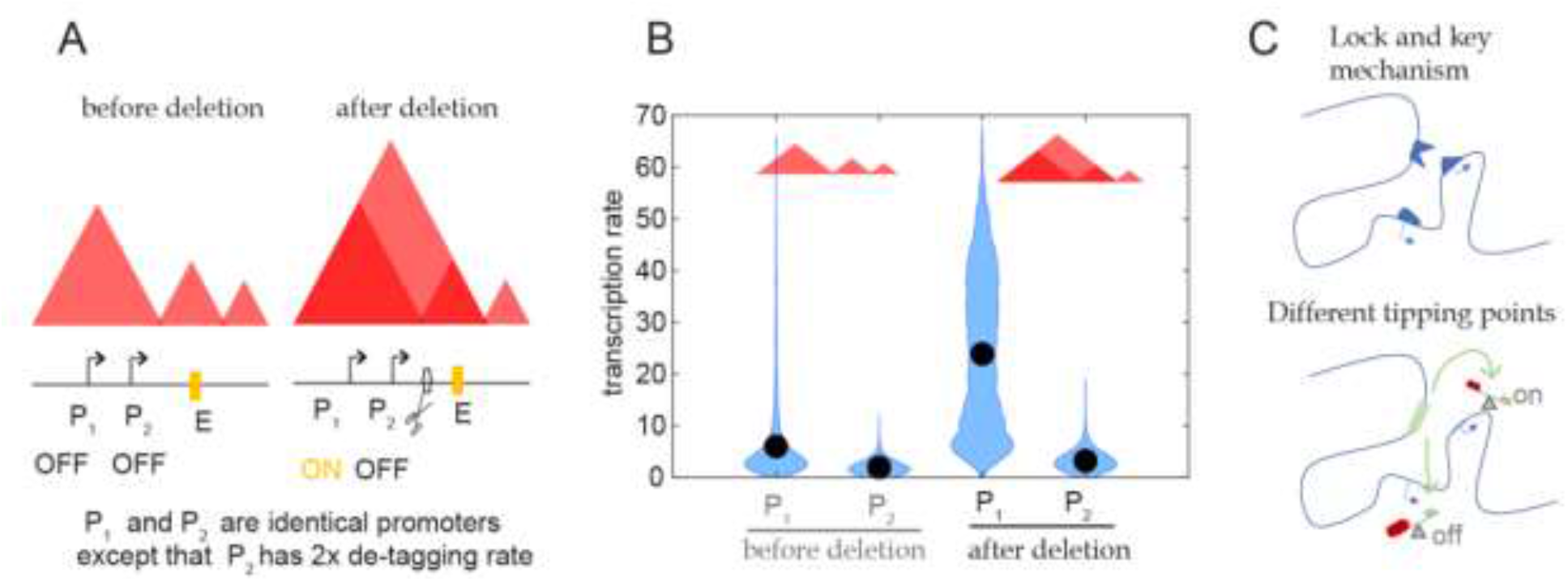
Differential sensitivity to E-P contact can lead to an illusion of specificity. **(A)** Model setup, depicted schematically. Two simulated promoters undergo the same change in enhancer contact frequency after a TAD-border deletion. **(B)** Violin plots of the changes in transcription rates (equivalently, promoter tag levels) after a two fold increase in the E-P contact rate for each promoter, simulating the border deletion depicted in A. **(C)** Cartoon representation contrasting the “lock and key” explanation of enhancer-promoter specificity with the “tipping point” explanation suggested by the results in B.

### Hysteretic in futile-cycle promoters creates memory at multiple timescales, a potential explanation why global TAD disruption causes little transcriptional change

We next examined the population level transcription dynamics of simulated cells exposed to a mild change in E-P contact, like the 2-fold or less expected from TAD border disruption due to border deletion or cohesin removal (**Fig. 3A & B**). We focused this analysis on promoters in the hypersensitive regime, since promoters in the sub-linear regimes exhibit limited sensitivity to mild changes in E-P-contact. Simulated cells with a promoter started in an active state which experienced a two-fold reduction in contact frequency showed little change in expression by the end of the early equilibration period (**Fig. 3C**). However, by the late time point, the most cells exhibited dramatically decreased expression (**Fig. 3C**). This response was much slower than observed for inactivation of an enhancer, where most cells dropped to low expression during the early period (**Fig. 3C**). Thus, the transcription output of these promoters is robust to short structural perturbations, even though it remains rapidly responsive to chemical signalling changes, such as deactivation of an enhancer.

**Figure 3.**
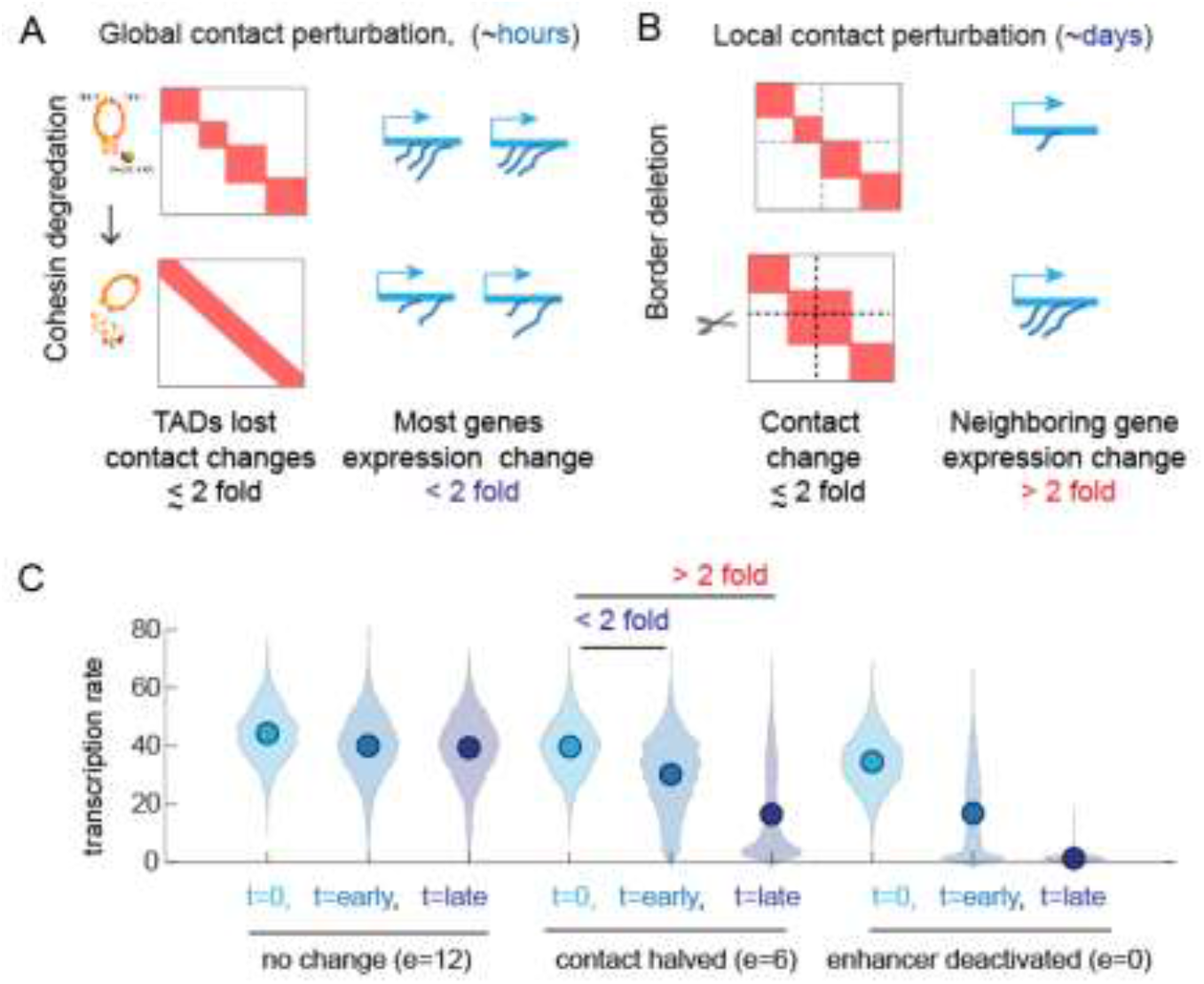
Transcriptional effects of contact perturbation depend on experiment timescale. **(A)** Schematic of global cohesin loop disruption experiments. **(B)** Schematic of individual TAD boundary deletion experiments. **(C)** Simulation of futile cycle promoter transcription rate as function of time under different enhancer perturbations.

These very different transcriptional behaviors in response to the same perturbation resemble recently reported experimental results. Global disruption of TADs and loops results in only subtle changes in transcription (Rao et al., 2017), yet disruption of individual TADs by deleting boundary sequences often resulted in significant expression changes to nearby genes (**Fig. 3A vs. B**) (reviewed in: (McCord et al., 2020), see also **Supp. Fig. 4**). Notably, global disruption was enabled by the rapid response of degron technology, and transcription assayed just hours later to avoid the complications of secondary effects from removing cohesin from the genome. In contrast, in local boundary deletion experiments, the requirement to clonally expand the initial mutant cell introduces multiple days and multiple cell-cycles of separation during which the effects of the lost contact stabilize. This suggests the 6-hour transient removal of cohesin by degrons may not be enough time to overcome the hysteretic nature of the promoter’s sensitivity to enhancer contact, inherent in the futile-cycle promoter model.

The hysteretic behavior of the model was further examined by comparing the response to a change in contact frequency as a function of initial state. Promoters that start at low tag/transcription levels follow different response curves from those that start high (**Fig. 4A**). This results from feedback where promoters which have a high level of the tag recruit more of the tag addition enzyme and thus require less interaction from the enhancer to counteract tag removal. Consequently, these promoters generally stayed highly transcribing even at EP contact levels that are insufficient to overcome tag removal from promoters that started with few tags. As illustrated in the simulations above, the degree difference between the curves depends on time (**Fig. 4B**). Promoters that experienced a very low rate of E-P contact rapidly converged to acommon distribution in which most of the population is silent, independent of the starting state. Those that experienced a high rate of contact rapidly converged to a distribution which is mostly actively transcribing. Promoters with intermediate contact rates exhibited much longer memory of the starting state (**Fig. 4B**).

**Figure 4.**
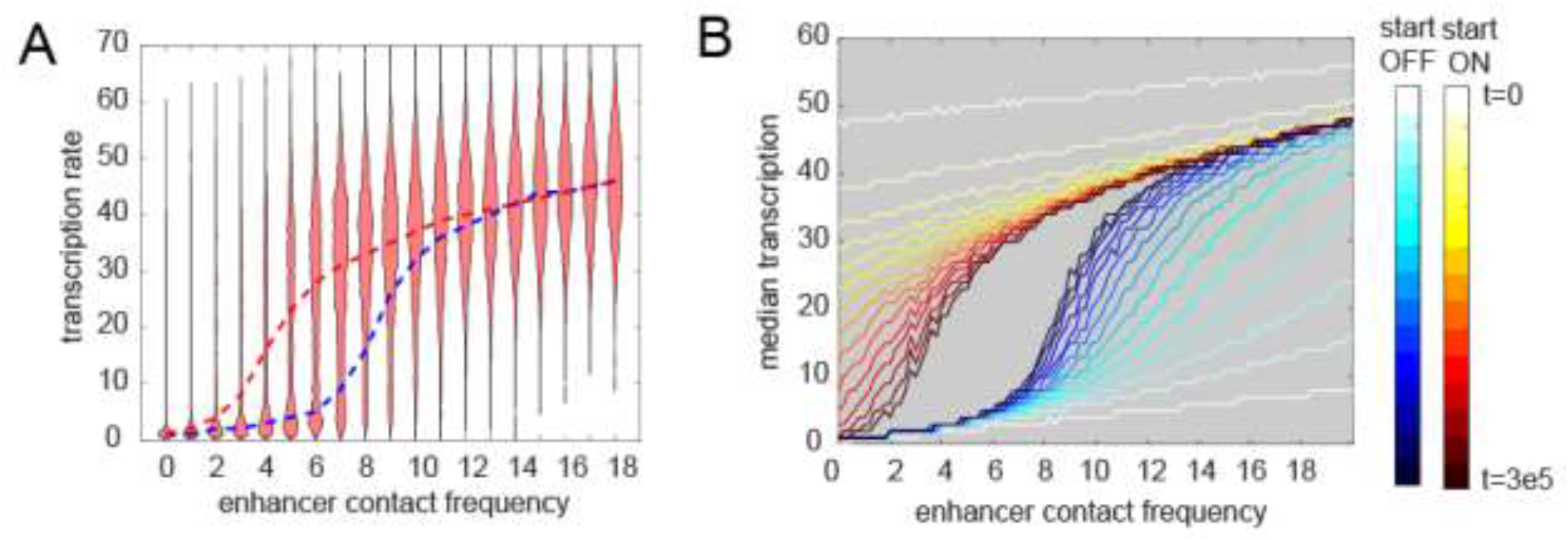
Transcription exhibits hysteresis with respect to enhancer contact frequency. **(A)** Violin plots of transcription as a function of enhancer contact, as in Fig 1D, but starting from a population in which promoters start in the highly-tag ‘ON’ state instead of the no-tag ‘OFF state’. Red dashed line connects the median behavior from this simulation. Blue dashed line shows median behavior starting from the ‘OFF’ state for comparison. **(B)** Median transcription rate for populations starting in the ‘ON’ or ‘OFF’ state and observed at different times after initialization of the simulation as indicated by the color legend.

### Transcription and enhancer-promoter contact may be largely uncorrelated in single cells

Having examined the temporal dynamics at the population scale, we turned the dynamic behaviors in single cells. We asked whether a promoter driven by contact-dependent enhancer activity and futile cycles, as described above, would be expected to show strong or weak correlation between E-P looping and transcription at the single cell level. This parallels experimental measurements we and others recently made (**Fig. 5A & B**). From our simulated cells, we computed the odds ratios for observing transcription in a window given observation of contact in that window (**Fig 5C**). Across the different regimes of E-P contact frequency, the Odds Ratios are statistically greater than 1, but only by a small margin of 0-30%, paralleling our recent empirical observation (Mateo et al., 2019). Since the frequency of interaction only tips the scales in the existing futile cycles of promoter modification, rather than acting as a triggering event for transcription, a strong correlation is not expected. The Odds Ratio still remains different from 1 however, as promoters which are experiencing a high frequency of EP contact are more likely to transcribe, increasing the probability the two events co-occur.

**Figure 5.**
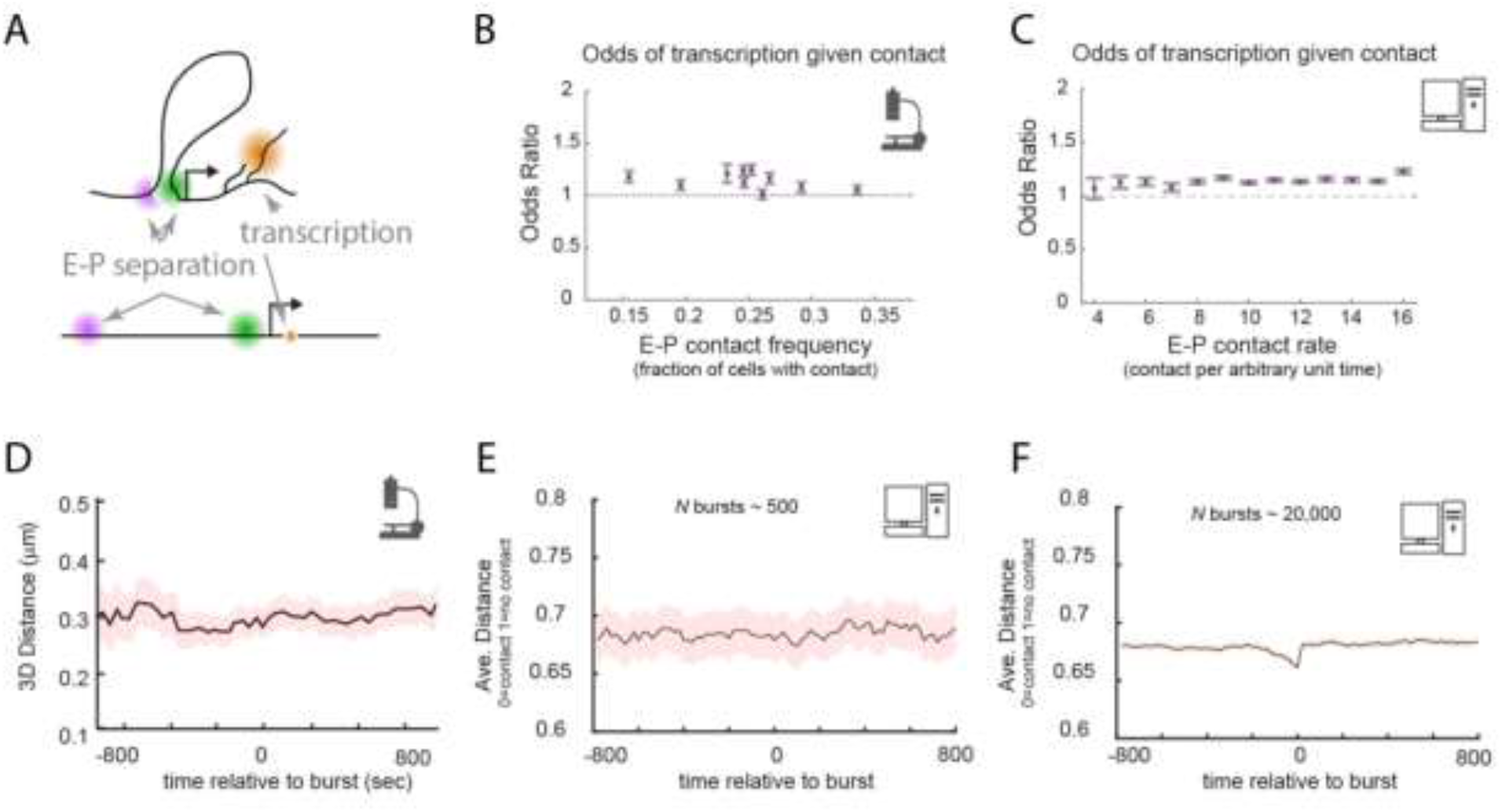
E-P contact events may exhibit minimal correlation to transcription events in single cells. **(A)** Schematic of the text-book expectation of correlation between E-P contact and transcription. **(B)** Odds ratios for observing nascent transcription given E-P proximity in recent super-resolution imaging experiments, sorted by observed E-P contact frequency. Experimental data from (Mateo et al., 2019). Error-bars show standard error from bootstrapping. **(C)** As in (B), but from simulations of futile cycle promoters across a range of E-P contact rates. **(D)** Average measured distance between enhancer and promoter probes relative to the time of transcriptional bursts, reproduced from (Alexander et al., 2019). **(E)** Average distance from simulations of futile cycle promoters, where 0 is contact and 1 is no contact, relative to time transcription burst, modeled as a tag concentration > 20. Shaded region denotes +/− standard error of the mean. **(F)** As in E, with sample size increased to 40 fold.

To further study the relation between E-P contact and transcription, we examined contact frequency in the stochastic model relative to the timing of transcription events. Previously, the lack of change in E-P distance across any time scale relative to detected transcription bursts was interpreted to refute a contact-dependent model of gene enhancer-mediated regulation (Alexander et al., 2019) (**Fig. 5D**). For a moderate number of simulated bursts (~500), we also find no detectable increase in proximity at any timescales relative to that of the burst (**Fig. 5E**). This is surprising at first, since by construction transcription is dependent on contact. Because the transition also depends on promoter-intrinsic tag addition and removal events, which also occur stochastically, many E-P contact events don’t result in transcription and many that do still exhibit variable timing, so that with small numbers of total events the dependence is washed out in the stochastic noise. Simulations involving a similar number of total cells as experiments also lacked detectable correlation between the fraction of time spent transcribing and average E-P distance or contact rate for individual cells, same as observed in the experimental data (**Supp Fig 5 A,B,D,E**). Substantially increasing the number of simulated cells uncovered weak but statistically significant correlations in each of these cases (**Fig. 5E**, and **Supp. Fig. 5 C,F**), similar to the observations from the *Drosophila* data (**Fig 5A**). Thus, the Futile Cycle transcription model shows that enhancers could regulate genes like Sox2 or the Bithorax genes in a purely contact-dependent manner of communication, even without strong correlation between contact events and transcription arising at the single cell level.

### Minimal parameter requirements for hypersensitivity

Thus far we have examined the behavior of a simple, discrete chemical model of promoter regulation (**Fig. 1C**) in a regime in which the model exhibits hypersensitivity and hysteresis, using stochastic simulations. However, not all parameter values for tag addition rates and removal rates will support these behaviors. We used a theoretical systems analysis approach to determine general requirements on the parameter values needed for these dynamic properties to arise. From a simple mass-action analysis of the chemical system, we derive a closed-form expression for the rate of change in tag concentration (d[a]/dt) as a function of the current tag concentration ([a]) (**Fig. 6A**) (see Methods), also called a *phase portrait* (Strogatz, 2018). For an appropriate region of parameter space (discussed below), d[a]/dt has up to three unchanging values, or *critical points,* where the curve crosses zero (**Fig. 6D**). To the left of the first zero, d[a]/dt is positive, so small values of [a] will increase. To the right, d[a]/dt is negative, so [a] will decrease, meaning that this first zero is a stable point—small excursions around this point will return. By the same logic it can be seen the second zero is unstable, and the third is stable. This bistability produces hysteresis in the system: promoters which start with low levels of expression (or tag) slowly increase [a] as the E-P frequency is raised to the point where the left and center critical point merge (a *saddle-node bifurcation),* which leads to a sharp discontinuous jump in the stable expression levels to the high state (**Fig. 6B**).

**Figure 6.**
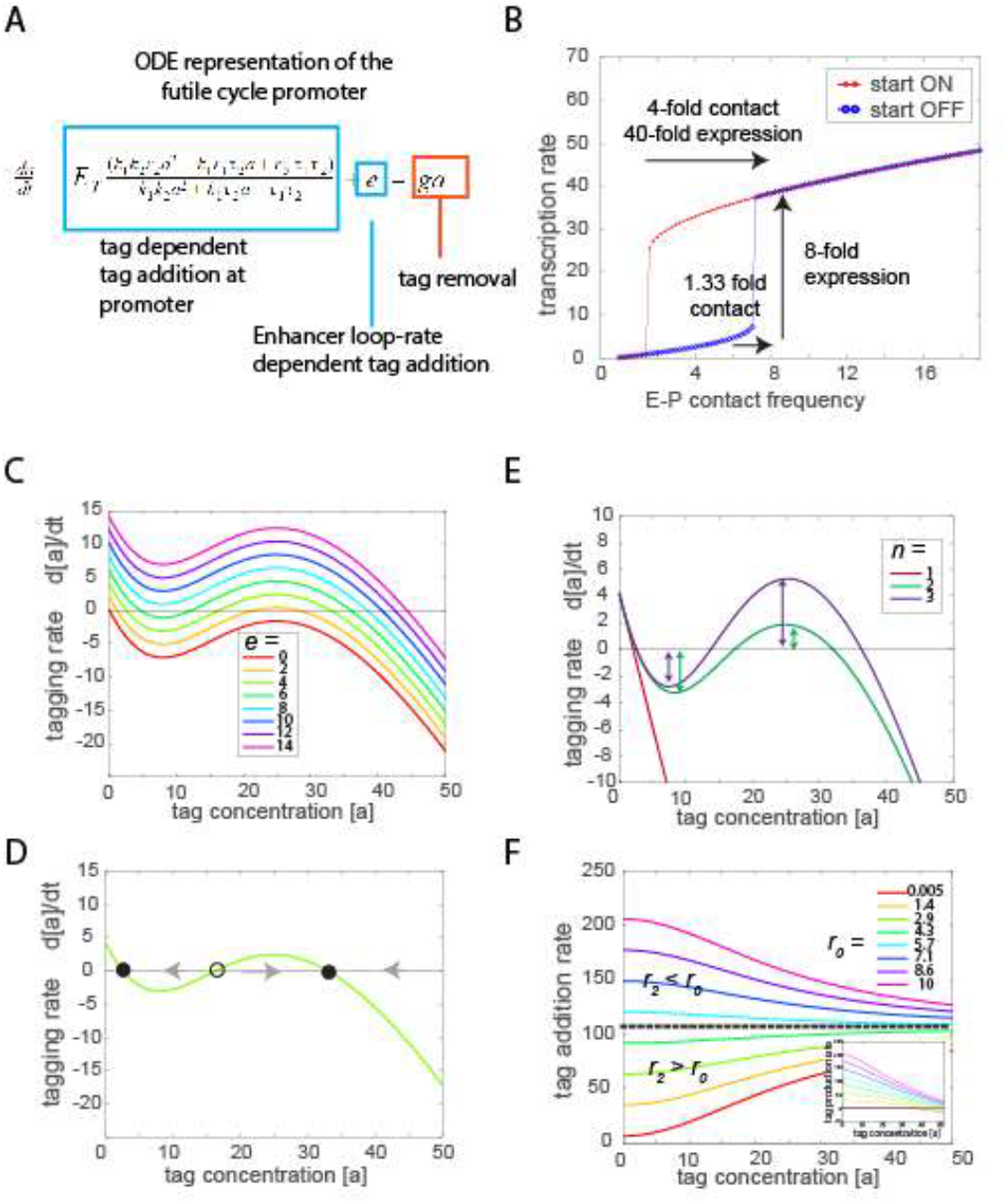
Phase portrait analysis of the futile cycle model. **(A)** Ordinary differential equation model for the Futile cycle promoter in the continuum limit, see Methods for derivation. **(B)** Transcription rate vs. E-P contact frequency for select parameters, illustrating the futile cycle supports hypersensitivity of transcription to E-P contact frequency and hysteresis. **(C)** Phase portrait illustrating how changing enhancer contact frequency shifts the nullcline and alters the number of stable-solutions (zero-crossing). **(D)** Phase portrait illustrating for the appropriate enhancer contact frequency, the futile cycle has up to two stable fixed points (solid circles) and one unstable fixed point (open circle). **(E)** Phase portrait showing that bistability arises for systems with at least two stimulation states. Increasing *n* > 2 produces a larger bistable regime. **(F)** Tag addition rate vs. tag concentration, illustrating the requirement for r2 > r0 needed for hypersensitivity and hysteresis.

We find two simple prerequisites for the Futile Cycle to exhibit bistable behavior. First, there must be at least three degrees of stimulation (*n_max_* ≥ 2) for the enzyme’s tag addition rate as a function of tag amount to be a sigmoidal curve, enabling three critical points (**Fig. 6E**) (see Supplementary Information for a derivation). Second, it requires *increasing* stimulation of the successive states (i.e. *r_n_* > *r*_*n*-1_), creating positive feedback in tag addition. If tag-associated states have less activity than non-associated states the production rate will not be an increasing function of total tag amount, preventing the existence of multiple stable states (**Fig. 6F**). These two requirements constitute all that is necessary for the general Futile Cycle model to exhibit hysteresis.

## Discussion

### Paradoxes and apparent contradictions arising from recent experimental results are reconciled by a minimal model of promoter activity

Here, we set out to understand one paradox, by what molecular mechanisms, if any, could subtle changes in chromatin structure result in large changes in transcription. We found that a simple model of promoter activity, inspired by the futile cycles of cell signalling, could readily exhibit the necessary hypersensitive response to changes in contact. Surprisingly, we also found properties of this model provide an unexpected explanation of three recent, controversial results. Here we discuss these three controversies, identify which features of the model offer explanation of the controversy, and identify experiments which may test these features in the future.

#### 1. Some promoters may be insensitive to contact perturbations, but still activated in a contact-dependent manner

An important obstacle to the hypothesis that 3D chromatin structures, including TADs, are important contributors to regulatory specificity of gene expression is the observation that many TADs harbor genes whose spatial-temporal patterns of expression differ substantially across development (Kvon et al., 2014; Schwarzer and Spitz, 2014; Symmons et al., 2014). For these patterns to differ, enhancers within the TAD must be able to regulate some promoters and not others even though all may experience similar frequency of EP interaction. A simple and common explanation of these differences is to conclude that contact frequency differences don’t play a role in regulation (since two promoters seeing similar contacts respond the same). Similarly, many enhancers are known to “hop over” intervening promoters (van Arensbergen et al., 2014)—activating expression of more distal genes without notably affecting the transcription of more proximal ones, such as Shh limb enhancer (Lettice et al., 2003) and several FGF8 limb enhancers (Marinić et al., 2013) or the snail shadow enhancer in flies (Perry et al., 2010). In most of these cases, the more distal gene experiences less frequent contact in available measurements. Collectively, these observations have been invoked to hypothesize that kilobase-to-megabase scale changes in chromatin structure (differences in contact frequency) may have little importance to regulatory specificity, and to suggest other mechanisms, such as combinatorial codes of transcription factors that create distinct lock-and-key recognition, may be the primary features contributing specificity to the genetic code (Ghavi-Helm et al., 2019; Kumar et al., 2020; Mir et al., 2019).

Our simulations indicate such conclusions are premature. A futile cycle promoter exhibits not just hypersensitive response to contact interactions, but also substantial regimes of sublinear or insensitive response. Promoter specific differences in the intrinsic addition or removal of tags can render one promoter unresponsive to an enhancer that activates a second promoter, even though the two experience similar absolute interaction frequency. Indeed, the unresponsive one may even experience a higher interaction frequency (e.g. because it is physically closer). It is thus premature to interpret such results as evidence of biochemical specificity, or to dismiss a role for chromatin structure. This is notable, since transgene assays indicate most regulatory elements will promiscuously activate most promoters if placed close enough (Arnold et al., 2013; Furlong and Levine, 2018; Kvon, 2015; Kvon et al., 2014). The model also suggests further experiments to test this mechanism. In particular, promoters which exhibit hypersensitive responses to certain perturbations, such as increased contact upon insulator removal, should be relatively insensitive in other regimes, which could be tested by secondary mutations. For example, further increasing in the contact between *Ubx* enhancers and the *abd-A* promoter following deletion of the border element, e.g. by removing some of the 30-60 kb which still separates them, is predicted to have primarily sublinear effects on promoter activity. Promoters which are insensitive to mild alterations to contact frequency, such as disruption of TAD boundaries, should still respond to larger changes in contact frequency, such as fusion of an enhancer immediately up or downstream. For example, the disruption of a single CTCF binding site at the border of the TAD containing Shh was recently reported to result in moderate structural change but no detected expression change (Williamson et al., 2019). If the Shh promoter is in the right parameter regime, the model predicts a further, similarly moderate structural change upon deletion of both CTCF sites at this border will create measurable change.

#### 2. Understanding global disruption of TADs

Examination of the time sensitivity of the model provided a possible explanation of a second set of controversial experiments. It was reported in 2017 that rapid removal of cohesin from the genome substantially altered global contact patterns, removing TADs and so-called “loops” or corner points from contact maps genome wide (Rao et al., 2017). Surprisingly, this structural perturbation resulted in little transcriptional changes, hardly any gene changing nascent transcription levels by more than a factor of two. This apparently contradicted results in earlier and concurrent reports, in which disruption of single TAD boundaries resulted in substantial change in expression to the local genes, leading to recent debate in the community. While it is important to note that the experiments were conducted in different cellular backgrounds, our modeling work demonstrates that a simple explanation is plausible. Two fold or smaller differences in contact frequency in the model show significant hysteresis, and will retain their initial transcription rate for many hours after such perturbation, *but* this memory is lost on longer time scales. As transcription was assayed hours after the contact changes in the degron experiment, little change was detected, whereas in experiments which genetically deleted border elements, transcription was assayed days and cell generations later, at which point larger effects were detected for some promoters.

Notably, TAD boundaries have been globally weakened or removed through a variety of alternative methods, including degradation of CTCF, deletion of the Cohesin associated factor NIPBL, protease cleavage of Cohesin, and siRNA (Nora et al., 2017; Schwarzer et al., 2017; Zuin et al., 2014). A direct comparison between the studies is complicated by the variations in cell type and genetic background used. Yet, by and large, transcriptional assays performed at longer intervals after first detection of E-P contact disruption exhibit greater transcriptional changes, consistent with the explanation suggested by these simulations.

How can we connect modeling time scales with the experimental measurements? While too little is known about the detailed chemical kinetics in the *in vivo* context to confidently extrapolate these results to absolute time scales, it is clear there is both a fast and a slow regime, with moderate change in tagging rates (or looping rates) associated with a slower time scale. Available *in vivo* data on recruiting epigenetic modifiers to promoters predominately shows significant response on the time scale of hours (Braun et al., 2017; Hathaway et al., 2012; Ho et al., 1996), but subtle responses can be detected with stronger recruitment in as little as a quarter hour in some studies (Braun et al., 2017). If promoter tags of our model are realized by such epigenetic mechanisms (discussed below), our results are consistent with the expectation that a decrease or increased contact frequency over only 6 hours will be insufficient to change the promoter state, while a substantially longer period such as the multiple days/generations between deletion and measurement can give the promoter enough time to change states.

#### 3. Transcription may be dependent on E-P contact even when the two aren’t correlated in single cells

It has been widely expected that active promoters should colocalize with their active enhancers, based largely on bulk population assays such 3C and H3K27ac Hi-Chip (Bartman et al., 2016; Deng et al., 2012; Kagey et al., 2010; Mumbach et al., 2017, 2016; Rao et al., 2014; Williamson et al., 2016), for review see (Furlong and Levine, 2018). Surprisingly, recent single cell measurements found that cells with enhancer-promoter proximity were only slightly more likely to exhibit nascent transcription than those lacking contact (Finn and Misteli, 2019b; Mateo et al., 2019). It is possible the correlation between looping and expression is missed in these fixed cell assays because the loops are transient relative to the rate of transcription, such that loops dissociate by the time the nascent RNA transcript is detectable. However, recent live cell imaging experiments suggest otherwise: live-cell tagging of chromatin proximal to the enhancer and promoter of Sox2 coupled with live-cell measurements of nascent RNA transcription uncovered no significant correlation between E-P proximity and transcription activity (Alexander et al., 2019). These data have supported speculation that enhancers must be able to influence promoter activity from a distance, e.g. through bridging by large protein condensates (Alexander et al., 2019; Benabdallah et al., 2019; Heist et al., 2019).

The model analyzed here clearly demonstrates that it is possible for transcription to be regulated by enhancer-promoter contact and yet show little temporal correlation, even in single-cell time-lapse data. This is because the E-P contact only changes the promoter state, toggling between a condition in which transcription events are likely, and one in which they are less likely. This explanation can be directly tested with larger datasets—with sufficient observations, a weak correlation should be detectable (as seen in the fixed cell data), since cells experiencing a higher frequency of contact are more likely to have a high level of tag and thus more likely to transcribe, even though it is not a one-to-one dependency.

### Repression also supports hypersensitivity and hysteresis

While for simplicity the model assumed that the tags accumulated at the promoter are activating, such that transcription is proportional to tag level, it is equally plausible that futile cycles of repressive tags at the promoter account for hypersensitivity of some promoters. In this case we model transcription as inversely proportional to the concentration of the promoter tag. The distal regulatory element which adds more of the tag should be termed a silencer rather than an enhancer. In response to increased contact with the silencer, the promoter transitions through a weakly-sensitive ON regime, a hypersensitive bistable regime, to a weakly-sensitive OFF regime (**Supp. Fig. 3**). Alternatively, a model in which the enhancer removes the repressive tag from the promoter supports all three regimes (**Supp. Fig. 3**).

### Histone modifications and/or phase-separated TF aggregates exhibit the key properties of tags needed to support hypersensitivity and hysteresis

Histone modifications are an obvious candidate for tags which the model assumes are added and removed in futile cycles and determine the transcriptional potential of the promoter. Several examples have been shown to exhibit the necessary features of positive feedback and competition between tag writers and erasers that are co-expressed. The histone methylation tags, H3K27me3 and H3K9me3 are both associated with silent promoters. The enzyme complexes which deposit these tags, PRC1 and SuVar(3,9), also preferentially associate with them compared to unmodified histones (Kim and Kingston, 2020; Sanulli and J Narlikar, 2020; Schuettengruber et al., 2017). This read-write feedback will produce the product-stimulation, which is critical for hysteresis, by increasing the residence time of the enzyme at already modified chromatin. Additional complexes, such as PRC1 in the case of H3K27me3 and HP1 in the case of H3K9me3 tags, provide further opportunity for read-write feedback through homotypic association, tag binding, and recruitment of tag writers (Kim and Kingston, 2020; Sanulli and J Narlikar, 2020; Schuettengruber et al., 2017). Known demethylases for either of these modifications, such as UTX and KDM4 provide the necessary futile cycle between adding and removal of tags required in the model (Kim and Kingston, 2020; Sanulli and J Narlikar, 2020; Schuettengruber et al., 2017). Known enzymes and protein complexes associated with deposition of tags positively correlated with transcription, such as H3K4me3 and H3K27ac, similarly exhibit product stimulated feedback. For example the histone acetylase P300/CBP associates with the transcription factor BRD4, which binds acetylated histones through its bromodomain (Black et al., 2006; Sabari et al., 2018). Cognate demethylases and deacetylases complete the futile cycle (Piunti and Shilatifard, 2016; Schuettengruber et al., 2017).

Notably, the Futile Cycle model as discussed here doesn’t require the “tag” be a histone modification or even an enzyme product. Any molecular group which can accumulate at the promoter and exhibit a higher likelihood of molecular addition to promoters which already contain the molecule will do. Micro-condensates of transcription factors are another interesting candidate. Individual molecules fusing and re-dissolving out of the condensate provide the necessary features of a futile cycle. Larger condensates are more likely to retain molecules and grow larger, since they offer more weak multivalent interactions, a process called Ostwald Ripening (Marqusee and Ross, 1983), and are more likely to fuse with other clusters, providing the necessary positive feedback. Many of the transcription factors associated with gene regulation have recently been shown to exhibit these properties, forming visible condensates of many molecules and exhibiting rapid recovery after photo-bleaching indicative of futile cycling into and out of the cluster (Strom and Brangwynne, 2019). These include PolII itself (Cho et al., 2018; Cisse et al., 2013; Guo et al., 2019; Sabari et al., 2018) transcriptional activators like Mediator and BRD4 (Cho et al., 2018; Sabari et al., 2018), and repressors like HP1 and CBX (Kim and Kingston, 2020; Sanulli and J Narlikar, 2020). As we develop improved tools to measure the protein occupancy of individual promoters in single cells along with chromatin structure, it will be possible to test the dynamic interplay between contact and protein / modification accumulation more directly.

### Enhancer cooperativity/super-enhancer hubs cannot explain hypersensitivity

It has been suggested cooperative, hub-like interactions among multiple enhancers targeting the same promoter may explain the hypersensitivity of transcription to enhancer-contact frequency. This proposal draws its intuition from the superficial similarity between the hypersensitivity of transcription to contact and the hypersensitivity to transcription factor (TF) binding that can arise through cooperative interactions among TFs. The idea is that while the individual E-P contacts may be weak and transient, much like individual TF-DNA interactions, if they stabilize one another, the system can show hypersensitive transitions: A perturbation removing a weak TAD boundary could subtly increase the contact frequency for all enhancers in a super-enhancer cluster one side of the boundary with a promoter on the other side -- much like a subtle increase in TF concentration can the binding probability for each TF with its target site. If this subtle increase in interaction is sufficient for a second enhancer to join the promoter before the first one dissociates, the cooperative interactions between them make the connection last even longer and so the third joins, further stabilizing the complex and so forth. Thus a small initial change has led to a dramatic change in total contact duration, which in this view explains the strong transcriptional changes without invoking promoter properties. However, available data in which contact frequency was measured before and after structural perturbation fail to support this model – perturbations of initially weak boundaries result largely in weak changes in absolute contact frequency, not the strong changes predicted by formation or collapse of a cooperative hub (e.g. Fig 1b, Supp. Figs 3-4).

## Conclusion

We have described and analyzed a simple model of enhancer-contact-dependent promoter activity. In this view, the promoter is not only a middleman in the regulation of transcription. It does not only relay the activity state of the enhancer into transcriptional activity of its target gene, nor function as a simple amplifier of enhancer activity to set the levels of expression. Instead, through accumulation of local tags, the promoter is able to integrate its history of interaction with one or more regulatory elements and respond in striking nonlinear ways to changes in these signals. By ‘accumulation of local tags’ we postulate nothing more than that the promoter is epigenetically regulated (in the broadest sense of ‘epigenetic). In addition to its core sequence, the accumulation of other factors at the promoter, be they histone modifications or transcription factors, provides the necessary mechanism for the dynamical system to exhibit memory.

This view of the promoter as an epigenetic regulator and integrator of signals is not new, and does not posit the existence of any unknown molecular mechanisms. However, a minimal model that captures the key features of such promoters we have shown can exhibit a much more complex dependency on chromatin structure than previously acknowledged. These behaviors offer a simple explanation to controversies of major importance to chromatin structure and transcription community. These include (1) why TAD boundaries can be weak and why insulators only mildly affect contact frequency and yet be major regulators of transcription (2) why many genes thought to be responsive to distal enhancers are insensitive to some structural changes (3) why global disruption of structure and acute disruption of structure have been reported to have distinct effects on transcription, and (4) why physical distance between known distal enhancers and their cognate gene does not show obvious correlation with transcriptional state.

## Methods

### Stochastic simulations

Stochastic simulations were written in Matlab(™) 2020a. Executable source code to run these simulations is available here https://github.com/BoettigerLab/transcription-modeling. A detailed table of parameters values and parameter descriptions is provided at the start of each script. The associated figure is indicated in the filename and in the header of the corresponding script for each simulation.

In brief, these simulations model the chemical master equation in explicit time and discrete time steps, with discrete transition probabilities for the addition and removal of promoter tags. For simplicity, we equate promoter modification and transcription, reducing the total parameter space without loss of generality.

### Continuum limit calculations

The ordinary differential equation (ODE) model was derived from the discrete chemical master equation using mass-action kinetics as described below. The stimulated enzyme example presented below is only one implementation of the futile cycle model, but used here because it’s the simplest and most intuitive system that recapitulates the desired quantitative behavior.

#### Model parameter definitions

Consider a promoter-bound enzyme that attaches molecular tags to a promoter at some basal rate *r*_0_. This enzyme can also associate with *n* promoter-bound tags up to some number *n_max_*, which changes the tagging rate to *r_n_*. The enzyme-acetyl association rates *k_n_* are dependent on the current association state (e.g. cooperativity), while dissociation is assumed to be constant (for now, but can be generalized later). “Free” tags not associated with the enzyme but still promoter-bound are removed from the promoter with a rate constant *g.*

#### Reaction and rate equations

For illustrative purposes, we will start with the *n_max_* = 2 case (later, we’ll show this is sufficient to capture nonlinear behavior of interest). Then the enzyme has three “association states” (*n* = 0, 1, or 2):

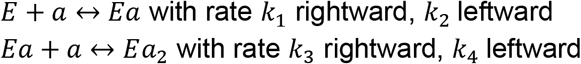

where *E* is enzyme and *a* is tag. In general, for *n* < *n_max_*:

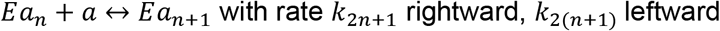

where *n* represents the association state.

#### Tagging/detagging

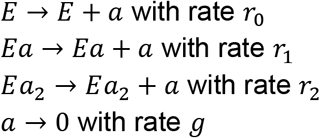

In general:

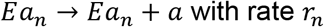

Furthermore, we assume that enhancer-promoter contact has the effect of adding more tags at the promoter such that

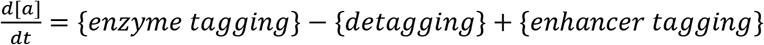

For now, assume a constant dissociation rate such that *k*_2_(_*n*+1_) = *τ,* and replace the subscripts 2*n* + 1 → *n* + 1 for easier notation. Then we have

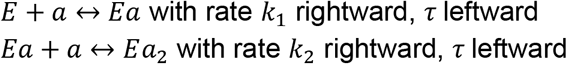

Now we can write out the rate equations:

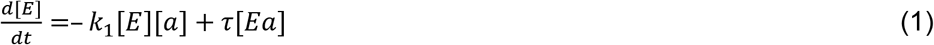

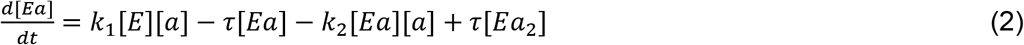

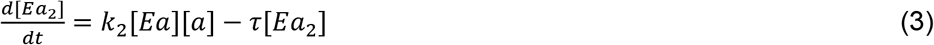

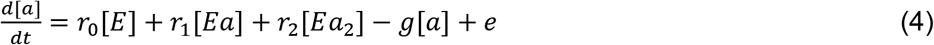

where *e* is the tagging contribution from enhancer-promoter contact. Assuming the total enzyme number is constant,

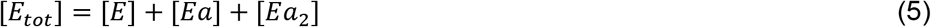

Our goal is to find 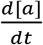 as a function of [*a*] and constants, and to study the steady state behavior of the system, i.e. finding the solutions for [*a*] when equations (1) through (4) are all set to zero. Solving (1) and (3), then substituting [*Ea*] and [*Ea_2_*] into (4) and (5):

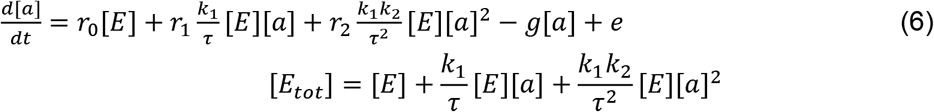

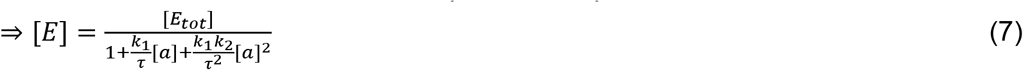

Combining (6) and (7):

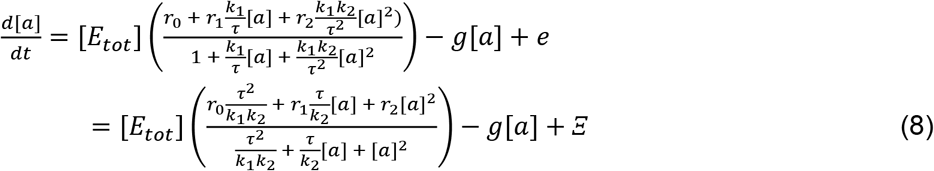

The terms with *r_n_* represent enzyme tagging, and the term with *g*is detagging. This nonlinear system with *n*=2 can be solved explicitly for the steady state values of *a*, though the solutions are less compact than the final equation, and are most easily interpreted in the phase portraits of equation 6. Analytic steady state solutions were calculated in Matlab using the symbolic algebra package. Scripts to reproduce these calculations are available on our github site at: https://github.com/BoettigerLab/transcription-modeling.

### Insulation Score Calculations

Several variations of this metric exist, but we define it as follows: first the contact-frequency map is normalized relative to linear genomic distance by dividing each off-diagonal row by the sum of the row. This improves the interpretability of the measure, as the inter-domain triangle contains more interactions further from the diagonal than the intra-domain ones (**Supp. Fig 3**). We then scan the whole genome, computing the average, normalized interaction-frequency within each of the 3 triangular regions shown in **Supp. Fig 3**: upstream (intra-domain U), downstream, intra-domain D, and between inter-window (domain I) of query point. We report the ratio of intra-domain to inter-domain interactions for this sliding window as the insulation score for that genomic position, as indicated in **Supp. Fig 3**. This metric can be computed for a range of sliding window sizes. As long as the window size is smaller than the typical TAD and large enough not to be sensitive to measurement-error fluctuations associated with over-binning the data, the distribution of insulation scores does not depend strongly on bin size. A comparison of several different computational algorithms for identifying TADs showed greatest agreement across replicates and robustness to data normalization using this metric (Gong et al., 2018; Lazaris et al., 2017).

Balanced, Hi-C matrices from (Rao et al., 2014) were downloaded and exported using Juicebox (Durand et al., 2016) for manipulation in Matlab for computing insulation scores.

## Acknowledgements

We thank Luca Giorgetti and Gregory Roth for informative discussions and members of the Boettiger lab for a critical reading of the manuscript. This work was supported by a CASI award from the Burroughs Wellcome Foundation and NIH New Innovator Award (DGM132935A) to A.N.B. J.X. was supported by the Molecular Biophysics Training Program at Stanford (GM008294). A.H. is supported by a Walter V. and Idun Berry Postdoctoral Fellowship.

## Supplemental Figures

**Supp. Figure 1.**
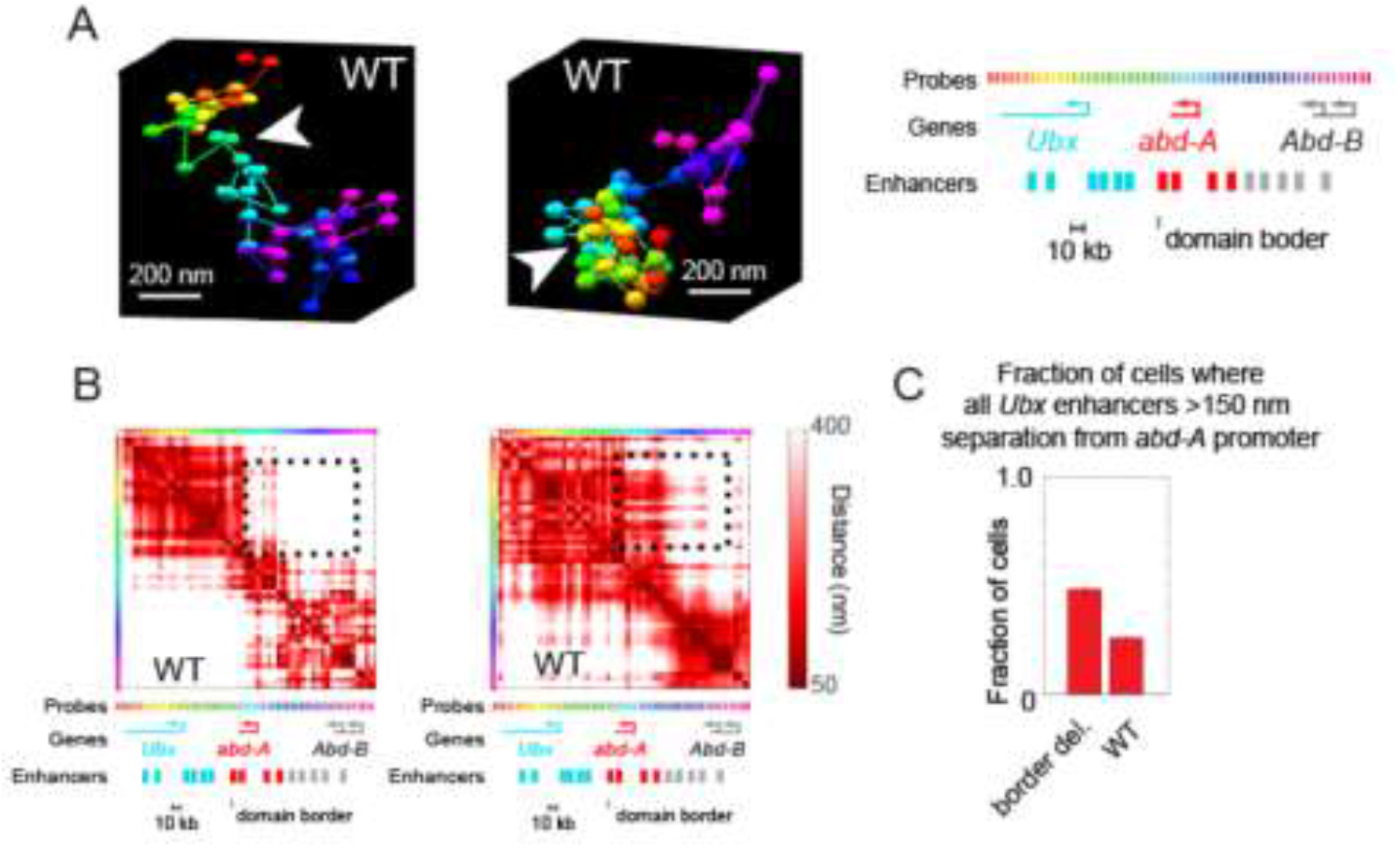
Wildtype cells also exhibit cross-border contacts. **(A)** Images of the Ubx and abd-A domain from two WT cells, reconstructed with Optical Reconstruction of Chromatin Architecture. White arrows mark the border between the domains, which are well separated in the first example and intermixed in the second. **(B)** Respective quantification of the pairwise distance between all labeled positions along the chromatin for the two cells shown in A. Black dotted box highlights the cross-border proximity found in the second cell. **(C)** Quantification of the frequency in which Ubx enhancers are in proximity to the abd-A promoter in WT vs. border deleted cells.

**Supp. Figure 2.**
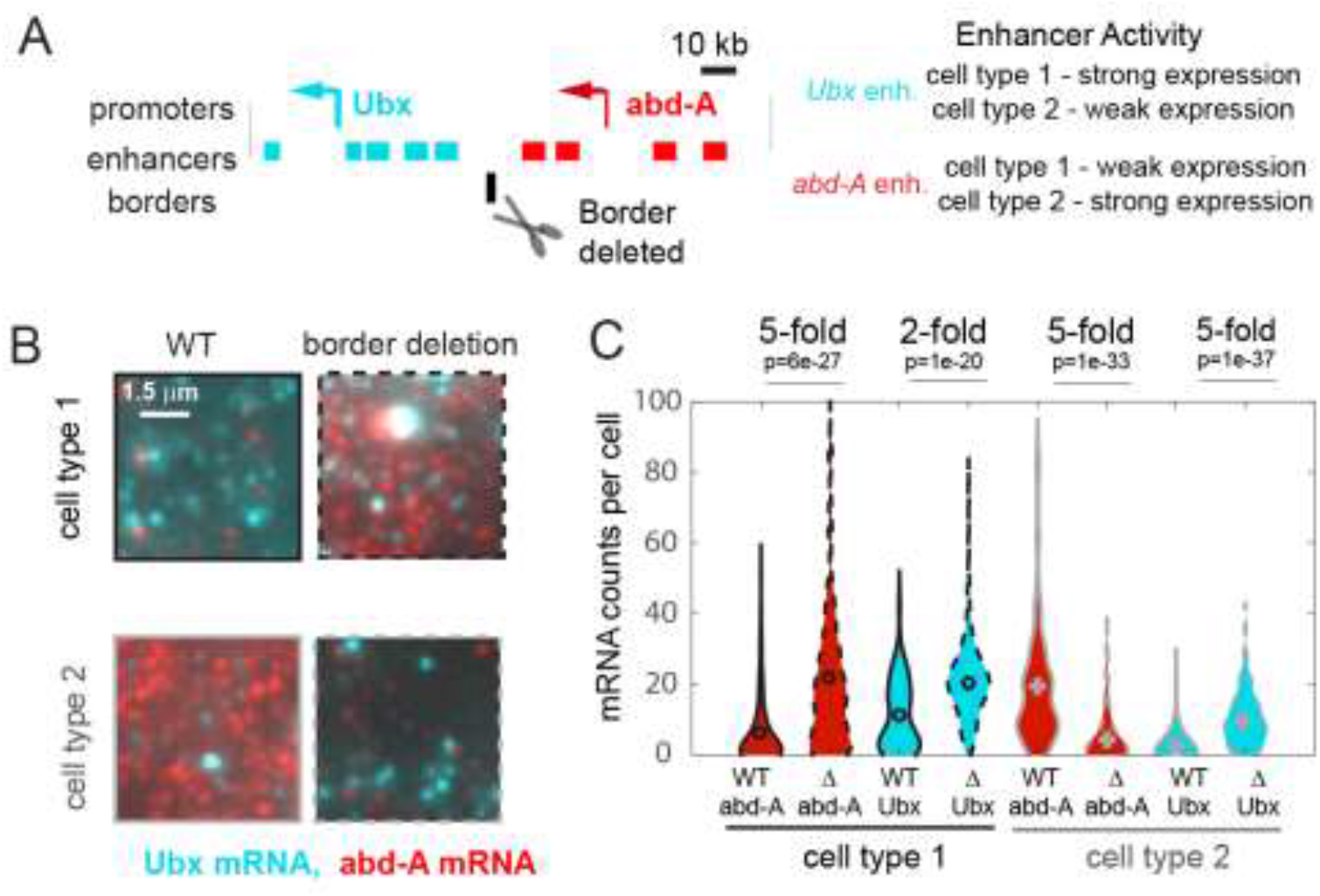
Quantitative effects of TAD border deletion on RNA expression. **A** Schematic of the genomic region containing the genes Ubx and abd-A and their surrounding enhancers. Both genes are expressed in two cell types of the abdominal region of the Drosophila embryo, under control of the indicated enhancers. Ubx enhancers drive strong expression in cell type 1 and weak expression in type 2 cells and v.v. for abd-A enhancers, as assayed by both individual enhancer-reporter transgenes and spatial analysis of the domain (Irvine et al., 1991; Mateo et al., 2019; Müller and Bienz, 1991; Qian et al., 1991). The position of the TAD border, removed by a 4kb deletion in ‘border deleted cells’ is also shown. **B** A zoom in view of smRNA FISH signal from single cells of either cell type, showing the typical difference in expression levels in wildtype cells and mutant cells. **C** Quantification of expression levels across the population, shown as a violin plot across all cells. The mRNA type (red = abd-A, cyan = Ubx), genotype (wildtype = solid outline, border-deletion = dotted outline), and cell type are indicated (black = type 1, grey = type 2), along with the median fold change in expression upon border deletion. Results from Wilcoxon comparison of median expression are given as *p*-values.

**Supp. Figure 3.**
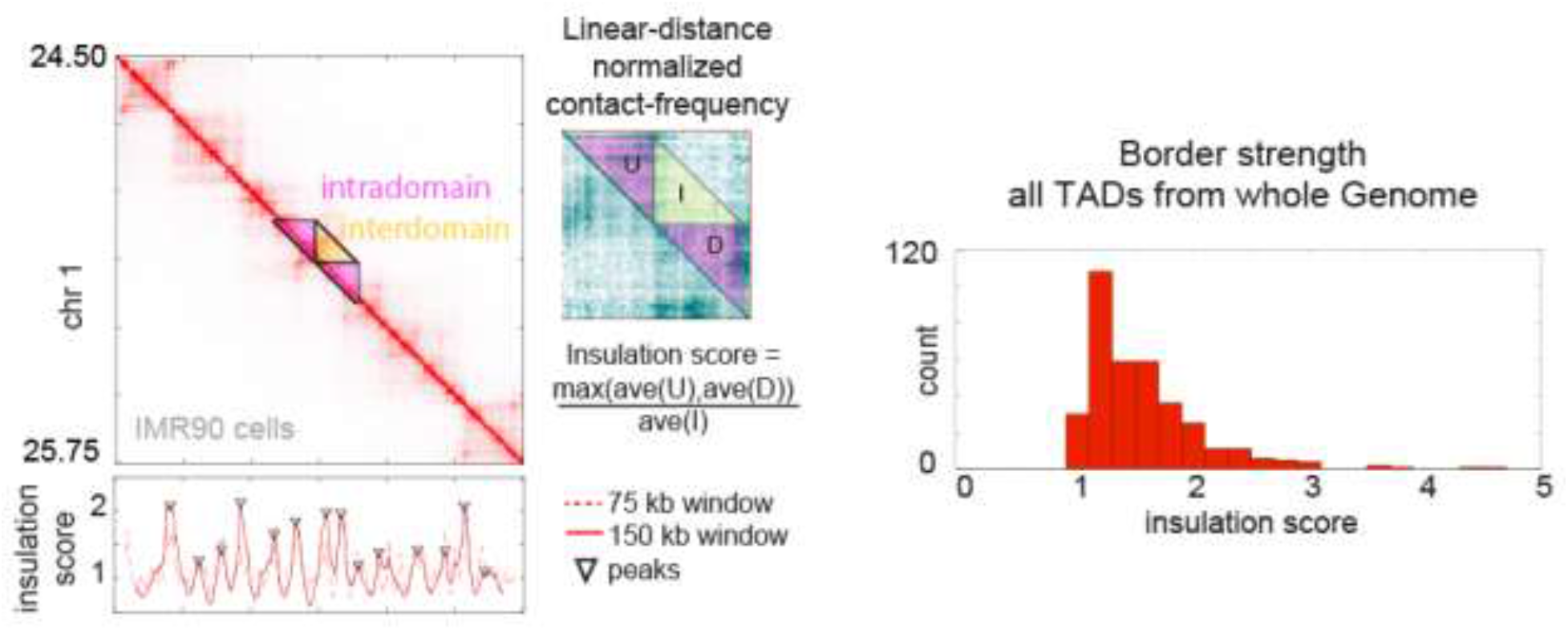
Calculation of TAD insulation scores across the genome using Hi-C data. Hi-C data from an arbitrary selected region of chromosome 1, exhibiting multiple TADs also known as contact domains. The blowout shows a zoom-in view of the masked region, following normalization to the linear-distance separation. The insulation score for this data computed with two different sliding window sizes is plotted below. Peak calls for the 150 kb window are shown, though both windows give similar results. Note these peaks correspond well to borders of the TADs seen by visual inspection. Right, a histogram of the insulation score for all peaks (TAD borders) called genome wide. Note, insulation score directly reflects the ratio of the preference for intra-TAD vs. inter-TAD contacts thanks to the prior normalization for linear distance effects (see Methods for more details).

**Supp. Figure 4.**
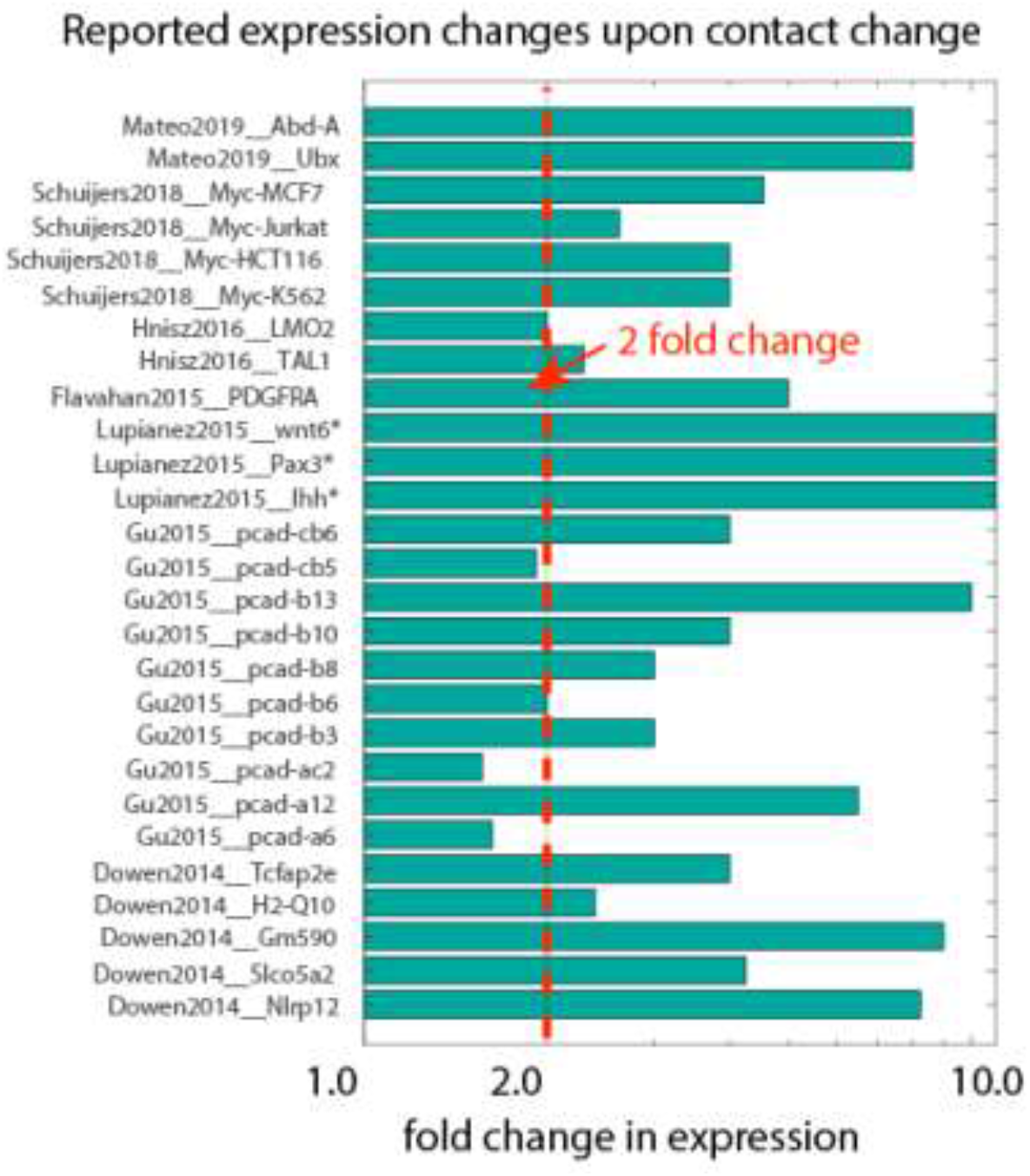
Compiled data on fold change in expression following deletion or inversion of TAD borders. Data are organized by source publication, indexed by first author and publication year and gene studied. In some cases expression was not appreciably detected prior to genomic perturbation, indicated by the “*”. Quantification techniques varied between experiments, and include smFISH, RNAseq, and qPcR. Dotted red line marks genes showing a greater than 2-fold change.

**Supp. Figure 5.**
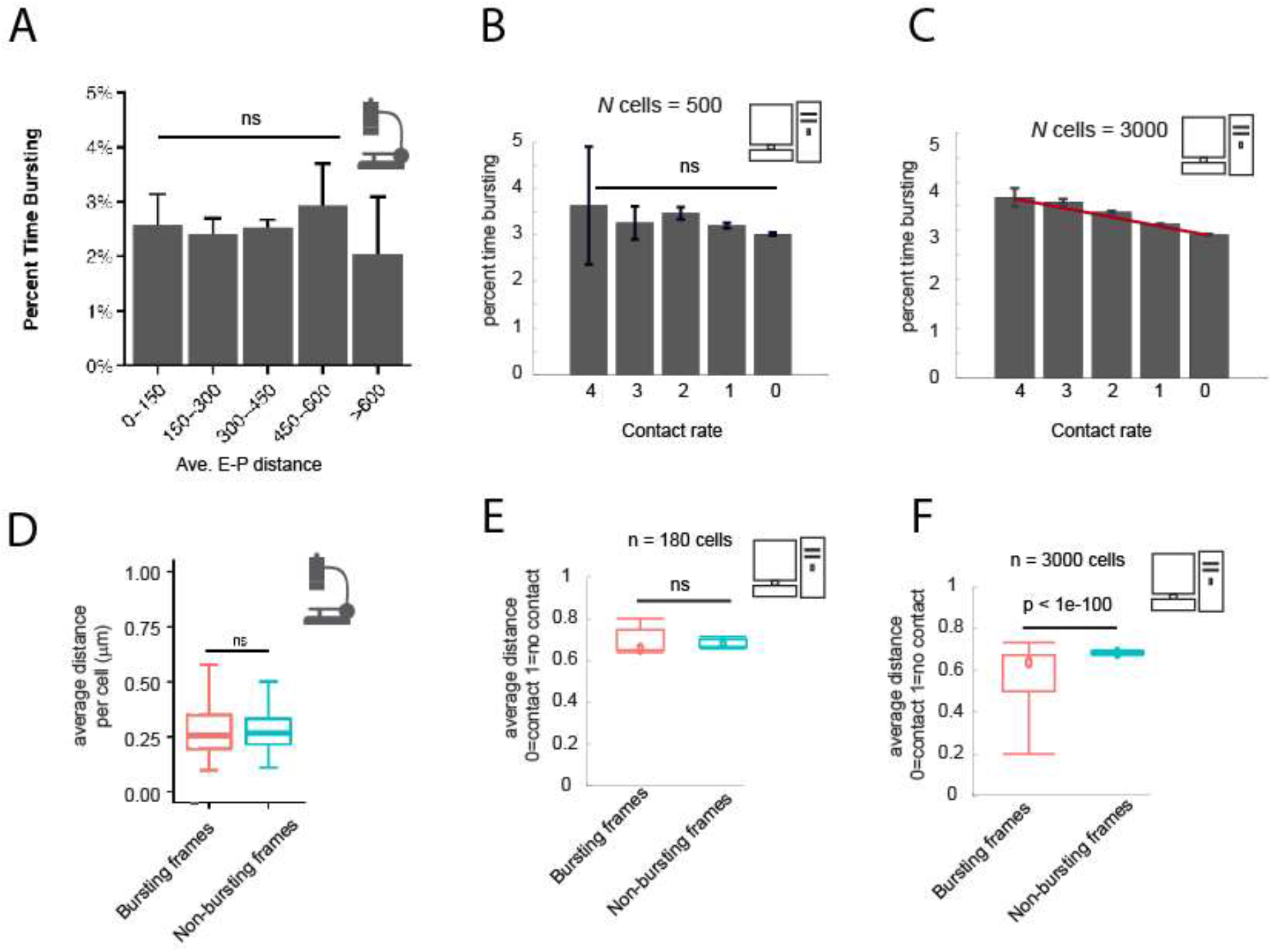
Comparison of experimental and simulation results on the correlation between E-P distance and time spent transcribing. **(A)** Data from (Alexander et al., 2019), quantifying the percent time bursting for cells binned by average measured distance between the enhancer and promoter probes. **(B)** Percent time in the bursting transcription state (promoter tag concentration > 20) for simulated cells, binned by their observed E-P contact frequency during the simulation window shown. N=500 cells. Error bars indicate 95% confidence limit around the median. The median time bursting for the high contact sample is not significantly different (*ns*) from any of the others by a Wilcoxon test (*p*>0.05). **(C)** As in (B), but with n=3000 cells, showing the emergence of a clear and statistically significant trend, Wilcoxon ranksum test, p<0.05. Errorbars indicate 95% confidence interval. **(D)** Box-plot analysis from (Alexander et al., 2019), showing no significant difference in the median distance between bursting and non-bursting promoters. **(E)**, as in **(D)** but with simulation data from the same simulations sampled in (B) Wilcoxon rank sum test results indicated by the bar. **(F)** as in (E) with n=3000 cells. Note, graph images in (A) and (D) are reproduced directly from (Alexander et al., 2019).

## References

Alberts B, Johnson A, Lewis J, Morgan D, Raff M, Keith Roberts PW, Others. 2018. Molecular biology of the cell.

Alexander JM, Guan J, Li B, Maliskova L, Song M, Shen Y, Huang B, Lomvardas S, Weiner OD. 2019. Live-cell imaging reveals enhancer-dependent Sox2 transcription in the absence of enhancer proximity. Elife 8. doi:10.7554/eLife.41769

Arnold CD, Gerlach D, Stelzer C, Boryń ŁM, Rath M, Stark A. 2013. Genome-wide quantitative enhancer activity maps identified by STARR-seq. Science 339:1074–1077.

Bartman CR, Hsu SC, Hsiung CC-S, Raj A, Blobel GA. 2016. Enhancer Regulation of Transcriptional Bursting Parameters Revealed by Forced Chromatin Looping. Mol Cell 62:237–247.

Beagrie RA, Scialdone A, Schueler M, Kraemer DCA, Chotalia M, Xie SQ, Barbieri M, de Santiago I, Lavitas L-M, Branco MR, Fraser J, Dostie J, Game L, Dillon N, Edwards PAW, Nicodemi M, Pombo A. 2017. Complex multi-enhancer contacts captured by genome architecture mapping. Nature. doi:10.1038/nature21411

Benabdallah NS, Williamson I, Illingworth RS, Kane L, Boyle S, Sengupta D, Grimes GR, Therizols P, Bickmore WA. 2019. Decreased Enhancer-Promoter Proximity Accompanying Enhancer Activation. Mol Cell. doi:10.1016/j.molcel.2019.07.038

Bintu B, Mateo LJ, Su J-H, Sinnott-Armstrong NA, Parker M, Kinrot S, Yamaya K, Boettiger AN, Zhuang X. 2018. Super-resolution chromatin tracing reveals domains and cooperative interactions in single cells. Science 362. doi:10.1126/science.aau1783

Black JC, Choi JE, Lombardo SR, Carey M. 2006. A mechanism for coordinating chromatin modification and preinitiation complex assembly. Mol Cell 23:809–818.

Braun SMG, Kirkland JG, Chory EJ, Husmann D, Calarco JP, Crabtree GR. 2017. Rapid and reversible epigenome editing by endogenous chromatin regulators. Nat Commun 8:560.

Cho W-K, Spille J-H, Hecht M, Lee C, Li C, Grube V, Cisse II. 2018. Mediator and RNA polymerase II clusters associate in transcription-dependent condensates. Science 4199:eaar4199.

Cisse II, Izeddin I, Causse SZ, Boudarene L, Senecal A, Muresan L, Dugast-Darzacq C, Hajj B, Dahan M, Darzacq X. 2013. Real-Time Dynamics of RNA Polymerase II Clustering in Live Human Cells. Science 664. doi:10.1126/science.1239053

Deng W, Lee J, Wang H, Miller J, Reik A, Gregory PD, Dean A, Blobel GA. 2012. Controlling Long-Range Genomic Interactions at a Native Locus by Targeted Tethering of a Looping Factor. Cell 149:1233–1244.

Dixon JR, Selvaraj S, Yue F, Kim A, Li Y, Shen Y, Hu M, Liu JS, Ren B. 2012. Topological domains in mammalian genomes identified by analysis of chromatin interactions. Nature 485:376–380.

Dowen JM, Fan ZP, Hnisz D, Ren G, Abraham BJ, Zhang LN, Weintraub AS, Schuijers J, Lee TI, Zhao K, Young RA. 2014. Control of Cell Identity Genes Occurs in Insulated Neighborhoods in Mammalian Chromosomes. Cell 159:374–387.

Durand NC, Robinson JT, Shamim MS, Machol I, Mesirov JP, Lander ES, Aiden EL, Durand NC, Shamim MS, Machol I, Rao SSP, Huntley MH, Aiden EL, Lieberman Aiden E, Berkum NL van, Williams L, Imakaev M, Ragoczy T, Telling A, Amit I, Lajoie BR, Sabo PJ, Dorschner MO, al. E, Paulsen J, Sandve GK, Gundersen S, Lien TG, Trengereid K, Hovig E, Stamenova EK, Bochkov ID, Robinson JT, Sanborn AL, Omer AD, Thorvaldsdóttir H, Winckler W, Guttman M, Getz G, Mesirov JP, Servant N, Nora EP, Giorgetti L, Chen C-J, Heard E, Dekker J, Barillot E, Zhou X, Lowdon RF, Li D, Lawson HA, Madden PAF, Costello JF, Wang T. 2016. Juicebox Provides a Visualization System for Hi-C Contact Maps with Unlimited Zoom. Cell Systems 3:99–101.

Finn EH, Misteli T. 2019a. A genome disconnect. Nat Genet 51:1205–1206.

Finn EH, Misteli T. 2019b. Molecular basis and biological function of variability in spatial genome organization. Science 365. doi:10.1126/science.aaw9498

Franke M, Ibrahim DM, Andrey G, Schwarzer W, Heinrich V, Schöpflin R, Kraft K, Kempfer R, Jerković I, Chan W-L, Spielmann M, Timmermann B, Wittler L, Kurth I, Cambiaso P, Zuffardi O, Houge G, Lambie L, Brancati F, Pombo A, Vingron M, Spitz F, Mundlos S. 2016. Formation of new chromatin domains determines pathogenicity of genomic duplications. Nature 1–15.

Furlong EEM, Levine M. 2018. Developmental enhancers and chromosome topology. Science 361:1341–1345.

Ghavi-Helm Y, Jankowski A, Meiers S, Viales RR, Korbel JO, Furlong EEM. 2019. Highly rearranged chromosomes reveal uncoupling between genome topology and gene expression. Nat Genet 51:1272–1282.

Gong Y, Lazaris C, Sakellaropoulos T, Lozano A, Kambadur P, Ntziachristos P, Aifantis I, Tsirigos A. 2018. Stratification of TAD boundaries reveals preferential insulation of super-enhancers by strong boundaries. Nat Commun 9:542.

Guo YE, Manteiga JC, Henninger JE, Sabari BR, Dall’Agnese A, Hannett NM, Spille J-H, Afeyan LK, Zamudio AV, Shrinivas K, Abraham BJ, Boija A, Decker T-M, Rimel JK, Fant CB, Lee TI, Cisse II, Sharp PA, Taatjes DJ, Young RA. 2019. Pol II phosphorylation regulates a switch between transcriptional and splicing condensates. Nature 572:543–548.

Guo Y, Xu Q, Canzio D, Shou J, Li J, Gorkin DU, Jung I, Wu H, Zhai Y, Tang Y, Lu Y, Wu Y, Jia Z, Li W, Zhang MQ, Ren B, Krainer ARR, Maniatis T, Wu Q. 2015. CRISPR Inversion of CTCF Sites Alters Genome Topology and Enhancer/Promoter Function. Cell 162:900–910.

Hathaway NA, Bell O, Hodges C, Miller EL, Neel DS, Crabtree GR. 2012. Dynamics and Memory of Heterochromatin in Living Cells. Cell 1–14.

Heist T, Fukaya T, Levine M. 2019. Large distances separate coregulated genes in living Drosophila embryos. Proc Natl Acad Sci U S A 116:15062–15067.

Hnisz D, Day DS, Young RA. 2016. Insulated Neighborhoods: Structural and Functional Units of Mammalian Gene Control. Cell 167:1188–1200.

Ho SN, Biggar SR, Spencer DM, Schreiber SL, Crabtree GR. 1996. Dimeric ligands define a role for transcriptional activation domains in reinitiation. Nature 382:822–826.

Irvine KD, Helfand SL, Hogness DS. 1991. The large upstream control region of the Drosophila homeotic gene Ultrabithorax. Development 111:407–424.

Jerković I, Szabo Q, Bantignies F, Cavalli G. 2020. Higher-Order Chromosomal Structures Mediate Genome Function. J Mol Biol 432:676–681.

Kagey MH, Newman JJ, Bilodeau S, Zhan Y, Orlando D a., van Berkum NL, Ebmeier CC, Goossens J, Rahl PB, Levine SS, Taatjes DJ, Dekker J, Young R a. 2010. Mediator and cohesin connect gene expression and chromatin architecture. Nature 2–8.

Kim J, Kingston RE. 2020. The CBX family of proteins in transcriptional repression and memory. J Biosci 45.

Kumar Y, Sengupta D, Bickmore W. 2020. Recent advances in the spatial organization of the mammalian genome. J Biosci 45.

Kvon EZ. 2015. Using transgenic reporter assays to functionally characterize enhancers in animals. Genomics 106:185–192.

Kvon EZ, Kazmar T, Stampfel G, Yáñez-Cuna JO, Pagani M, Schernhuber K, Dickson BJ, Stark A. 2014. Genome-scale functional characterization of Drosophila developmental enhancers in vivo. Nature 512:91–95.

Lazaris C, Kelly S, Ntziachristos P, Aifantis I, Tsirigos A. 2017. HiC-bench: comprehensive and reproducible Hi-C data analysis designed for parameter exploration and benchmarking. BMC Genomics 18:22.

Lettice LA, Heaney SJH, Purdie LA, Li L, de Beer P, Oostra BA, Goode D, Elgar G, Hill RE, de Graaff E. 2003. A long-range Shh enhancer regulates expression in the developing limb and fin and is associated with preaxial polydactyly. Hum Mol Genet 12:1725–1735.

Long HK, Prescott SL, Wysocka J. 2016. Ever-Changing Landscapes: Transcriptional Enhancers in Development and Evolution. Cell 167:1170–1187.

Lupiáñez DG, Kraft K, Heinrich V, Krawitz P, Brancati F, Klopocki E, Horn D, Kayserili H, Opitz JM, Laxova R, Santos-Simarro F, Gilbert-Dussardier B, Wittler L, Borschiwer M, Haas SA, Osterwalder M, Franke M, Timmermann B, Hecht J, Spielmann M, Visel A, Mundlos S. 2015. Disruptions of Topological Chromatin Domains Cause Pathogenic Rewiring of Gene-Enhancer Interactions. Cell 161:1012–1025.

Marinić M, Aktas T, Ruf S, Spitz F. 2013. An Integrated Holo-Enhancer Unit Defines Tissue and Gene Specificity of the Fgf8 Regulatory Landscape. Dev Cell 24:530–542.

Marqusee JA, Ross J. 1983. Kinetics of phase transitions: Theory of Ostwald ripening. J Chem Phys 79:373–378.

Mateo LJ, Murphy SE, Hafner A, Cinquini IS, Walker CA, Boettiger AN. 2019. Visualizing DNA folding and RNA in embryos at single-cell resolution. Nature 568:49–54.

McCord RP, Kaplan N, Giorgetti L. 2020. Chromosome Conformation Capture and Beyond: Toward an Integrative View of Chromosome Structure and Function. Mol Cell 77:688–708.

Mir M, Bickmore W, Furlong EEM, Narlikar G. 2019. Chromatin topology, condensates and gene regulation: shifting paradigms or just a phase? Development 146:dev182766.

Müller J, Bienz M. 1991. Long range repression conferring boundaries of Ultrabithorax expression in the Drosophila embryo. EMBO J 10:3147–3155.

Mumbach MR, Rubin AJ, Flynn RA, Dai C, Khavari PA, Greenleaf WJ, Chang HY. 2016. HiChIP: efficient and sensitive analysis of protein-directed genome architecture. Nat Methods 13:919–922.

Mumbach MR, Satpathy AT, Boyle EA, Dai C, Gowen BG, Cho SW, Nguyen ML, Rubin AJ, Granja JM, Kazane KR, Wei Y, Nguyen T, Greenside PG, Corces MR, Tycko J, Simeonov DR, Suliman N, Li R, Xu J, Flynn RA, Kundaje A, Khavari PA, Marson A, Corn JE, Quertermous T, Greenleaf WJ, Chang HY. 2017. Enhancer connectome in primary human cells identifies target genes of disease-associated DNA elements. Nat Genet 49:1602–1612.

Nora EP, Goloborodko A, Valton A-L, Gibcus JH, Uebersohn A, Abdennur N, Dekker J, Mirny LA, Bruneau BG. 2017. Targeted Degradation of CTCF Decouples Local Insulation of Chromosome Domains from Genomic Compartmentalization. Cell 169:930–944.e22.

Nora EP, Lajoie BR, Schulz EG, Giorgetti L, Okamoto I, Servant N, Piolot T, van Berkum NL, Meisig J, Sedat J, Gribnau J, Barillot E, Blüthgen N, Dekker J, Heard E. 2012. Spatial partitioning of the regulatory landscape of the X-inactivation centre. Nature 485:381–385.

Paliou C, Guckelberger P, Schöpflin R, Heinrich V, Esposito A, Chiariello AM, Bianco S, Annunziatella C, Helmuth J, Haas S, Jerković I, Brieske N, Wittler L, Timmermann B, Nicodemi M, Vingron M, Mundlos S, Andrey G. 2019. Preformed chromatin topology assists transcriptional robustness of Shh during limb development. Proc Natl Acad Sci U S A 116:12390–12399.

Perry MW, Boettiger AN, Bothma JP, Levine M. 2010. Shadow enhancers foster robustness of Drosophila gastrulation. Curr Biol 20:1562–1567.

Piunti A, Shilatifard A. 2016. Epigenetic balance of gene expression by Polycomb and COMPASS families. Science 352:aad9780–aad9780.

Qian S, Capovilla M, Pirrotta V. 1991. The bx region enhancer, a distant cis-control element of the Drosophila Ubx gene and its regulation by hunchback and other segmentation genes. EMBO J 10:1415–1425.

Quinodoz SA, Ollikainen N, Tabak B, Palla A, Schmidt JM, Detmar E, Lai MM, Shishkin AA, Bhat P, Takei Y, Trinh V, Aznauryan E, Russell P, Cheng C, Jovanovic M, Chow A, Cai L, McDonel P, Garber M, Guttman M. 2018. Higher-Order Inter-chromosomal Hubs Shape 3D Genome Organization in the Nucleus. Cell 1–14.

Rao SSP, Huang S-C, Hilaire BGS, Engreitz J, Perez EM, Kieffer-Kwon K-R, Sanborn AL, Johnstone SE, Bascom GD, Bochkov ID, Huang X, Shamim MS, Shin J, Turner D, Ye Z, Omer AD, Robinson JT, Schlick T, Bernstein BE, Casellas R, Lander ES, Aiden EL. 2017. Cohesin Loss Eliminates All Loop Domains. Cell 171:305–320.e24.

Rao SSP, Huntley MH, Durand NC, Stamenova EK, Bochkov ID, Robinson JT, Sanborn AL, Machol I, Omer AD, Lander ES, Aiden EL. 2014. A 3D Map of the Human Genome at Kilobase Resolution Reveals Principles of Chromatin Looping. Cell 159:1665–1680.

Redolfi J, Zhan Y, Valdes-Quezada C, Kryzhanovska M, Guerreiro I, Iesmantavicius V, Pollex T, Grand RS, Mulugeta E, Kind J, Tiana G, Smallwood SA, de Laat W, Giorgetti L. 2019. DamC reveals principles of chromatin folding in vivo without crosslinking and ligation. Nat Struct Mol Biol 26:471–480.

Rowley MJ, Corces VG. 2018. Organizational principles of 3D genome architecture. Nat Rev Genet 19:789–800.

Sabari BR, Dall’Agnese A, Boija A, Klein IA, Coffey EL, Shrinivas K, Abraham BJ, Hannett NM, Zamudio AV, Manteiga JC, Li CH, Guo YE, Day DS, Schuijers J, Vasile E, Malik S, Hnisz D, Lee TI, Cisse II, Roeder RG, Sharp PA, Chakraborty AK, Young RA. 2018. Coactivator condensation at super-enhancers links phase separation and gene control. Science 3958:eaar3958.

Sanulli S, J Narlikar G. 2020. Liquid-like interactions in heterochromatin: Implications for mechanism and regulation. Curr Opin Cell Biol 64:90–96.

Schuettengruber B, Bourbon H-M, Di Croce L, Cavalli G. 2017. Genome Regulation by Polycomb and Trithorax: 70 Years and Counting. Cell 171:34–57.

Schwarzer W, Abdennur N, Goloborodko A, Pekowska A, Fudenberg G, Loe-Mie Y, Fonseca NA, Huber W, Haering C, Mirny LA, Spitz F. 2017. Two independent modes of chromatin organization revealed by cohesin removal. Nature 551:51–56.

Schwarzer W, Spitz F. 2014. The architecture of gene expression: integrating dispersed cis-regulatory modules into coherent regulatory domains. Curr Opin Genet Dev 27:74–82.

Spielmann M, Lupiáñez DG, Mundlos S. 2018. Structural variation in the 3D genome. Nat Rev Genet 7:85–97.

Strogatz SH. 2018. Nonlinear Dynamics and Chaos with Student Solutions Manual: With Applications to Physics, Biology, Chemistry, and Engineering, Second Edition. CRC Press.

Strom AR, Brangwynne CP. 2019. The liquid nucleome - phase transitions in the nucleus at a glance. J Cell Sci 132. doi:10.1242/jcs.235093

Symmons O, Uslu VV, Tsujimura T, Ruf S, Nassari S, Schwarzer W, Ettwiller L, Spitz F. 2014. Functional and topological characteristics of mammalian regulatory domains. Genome Res 24:390–400.

van Arensbergen J, van Steensel B, Bussemaker HJ. 2014. In search of the determinants of enhancer–promoter interaction specificity. Trends Cell Biol 1–8.

Williamson I, Kane L, Devenney PS, Flyamer IM, Anderson E, Kilanowski F, Hill RE, Bickmore WA, Lettice LA. 2019. Developmentally regulated Shh expression is robust to TAD perturbations. Development. doi:10.1242/dev.179523

Williamson I, Lettice LA, Hill RE, Bickmore WA. 2016. Shh and ZRS enhancer colocalisation is specific to the zone of polarising activity. Development 143:2994–3001.

Zuin J, Dixon JR, van der Reijden MIJ a., Ye Z, Kolovos P, Brouwer RWW, van de Corput MPC, van de Werken H, Knoch T a., van Ijcken WFJ, Grosveld FG, Ren B, Wendt KS. 2014. Cohesin and CTCF differentially affect chromatin architecture and gene expression in human cells. Proc Natl Acad Sci U S A 111:996–1001.

